# Cargo diffusion shortens single-kinesin runs at low viscous drag

**DOI:** 10.1101/391284

**Authors:** John O. Wilson, David A. Quint, Ajay Gopinathan, Jing Xu

**Affiliations:** Physics; Center for Cellular and Biomolecular Machines, University of California, Merced, California 95343, USA

**Author notes:** **Address correspondence to:** David A. Quint, Ajay Gopinathan, and Jing Xu.

## Abstract

Molecular motors are mechanoenzymes that actively drive long-range transport in cells. Thermal diffusion of the cargo can result in mechanical load on the motor carrying the cargo; the direction of this diffusion-based load is not correlated with motor motion. Recent single molecule-based experiments highlighted a strong asymmetric dependence of the run length of the single kinesin-1 motor on load direction, raising the intriguing possibility that thermal diffusion of the cargo may non-trivially influence the run length of the motor carrying the cargo. To test this possibility, here we employed Monte Carlo-based stochastic simulations to evaluate the transport of single-kinesin cargos over a large parameter space of physiologically relevant solution viscosities, cargo sizes, and motor velocities. Our simulations uncovered a previously unexplored, significant shortening effect of cargo diffusion on single-kinesin run length. This effect is non-monotonically influenced by viscous drag force on the cargo, which biases the effect of cargo diffusion toward the hindering direction. The non-monotonic variation of cargo run length with drag force is the direct result of the asymmetric response of kinesin’s run length to load direction. Our findings may be important for understanding the diverse characteristics of cargo transport, including run length, observed in living cells.

## Introduction

Molecular motors such as kinesin-1 are mechanoenzymes that drive long-range transport of cargos within living cells (Hirokawa and Noda, 2008; Vale, 2003). This transport process is challenging to accomplish because motors must overcome substantial thermal diffusion to maintain directional transport. Thermal diffusion encompasses the set of random, non-directional motions that result from thermal agitation (Einstein, 1905). Thermal diffusion plays important roles in a variety of biological processes, including early embryonic patterning (Gregor et al., 2005; Turing, 1990), cell signaling (McMurtrey, 2017), and metabolism (Rohde and Price, 2007). For motor-based transport, thermal diffusion can manifest as random motions of the motor or of the cargo. A recent investigation highlighted the significant effect of thermal diffusion of individual motor domains on single-kinesin function in vitro (Sozański et al., 2015). How thermal diffusion of the cargo influences motor-based transport, however, has not been explored.

The functions of molecular motors are affected by external force, or “load” (Milic et al., 2014; Schnitzer et al., 2000). Recent single-molecule investigations revealed an intriguing asymmetric dependence of the distance traveled by a single kinesin-1 (“run length”) on load direction (Milic et al., 2014). Perhaps counterintuitively, under the same amount of load, kinesin’s run length is significantly *shorter* when the load is oriented in the same (“assisting”) versus the opposite (“hindering”) direction of motor motion (Milic et al., 2014). Of note, sensitivity to assisting load was not observed for kinesin’s transport velocity, another important biophysical function of the motor; kinesin maintained its unloaded velocity when the load was oriented in the assisting direction (Milic et al., 2014; Schnitzer et al., 2000).

Thermal diffusion of the cargo can exert load on the motor (Kunwar et al., 2008). Importantly, because cargo diffusion is not correlated with motor motion (Beausang et al., 2007; Einstein, 1905; Mogilner et al., 2002), the direction of the load from cargo diffusion can assist or hinder motor motion. Given the recently identified asymmetric response of kinesin’s run length to load direction (Milic et al., 2014), we hypothesized that cargo diffusion may non-trivially influence the run length of the kinesin carrying that cargo.

Here we employed Monte Carlo-based stochastic simulations to numerically examine the effect of cargo diffusion on transport by a single kinesin. We carried out our simulations over a large parameter space that captures crucial transport characteristics in living cells, including variations in viscosity (Dix and Verkman, 1990; Kalwarczyk et al., 2011; Kuimova et al., 2009; Luby-Phelps et al., 1993; Margraves et al., 2011; Suhling et al., 2004), cargo size (Bakker et al., 1997; Casley-Smith, 1969; Keller et al., 2017; Margraves et al., 2011; Shubeita et al., 2008; Wiemerslage and Lee, 2016; Zhang et al., 1998), and motor velocity (Lorenz and Willard, 1978; Tytell et al., 1981). Associated with this large parameter space is an additional load on the motor: the viscous drag force that hinders cargo motion (Laidler and Meiser, 1982). Our simulations revealed that cargo diffusion significantly shortens single-kinesin run length, an effect that is non-monotonically tuned by viscous drag force on the cargo.

## Results

### Thermal diffusion of the cargo shortens single-kinesin run length at low viscosity

We used a previously developed Monte Carlo simulation (Kunwar et al., 2008) to examine the effect of cargo diffusion on kinesin run length (Methods). In this simulation, the motor steps directionally along the microtubule track, while its cargo undergoes randomly oriented thermal diffusion (Beausang et al., 2007; Einstein, 1905; Kunwar et al., 2008; Mogilner et al., 2002). The displacement between the motor and the cargo determines both the magnitude and the direction of force experienced by the motor. The effect of load on run length is modeled as the motor’s load-detachment kinetics, which describes the effect of load on the stochastic probability of the motor becoming detached from its microtubule track per unit time (“detachment rate”). Previously, this model included consideration of kinesin’s load-detachment kinetics under hindering load only and assumed that the motor’s detachment rate is unaffected by assisting load (Kunwar et al., 2008). In the current study, we extended the load-detachment kinetics of the simulated motor (Methods) to reflect recent experimental measurements of the motor’s detachment rate under assisting load (Milic et al., 2014).

We first examined the run length of single-kinesin cargos over a physiologically relevant range of solution viscosities (Dix and Verkman, 1990; Kalwarczyk et al., 2011; Kuimova et al., 2009; Luby-Phelps et al., 1993; Margraves et al., 2011; Suhling et al., 2004), while holding the cargo size and motor velocity constant at 0.5 μm in diameter and 0.8 μm/s when unloaded, respectively (Figure 1). These parameter choices are commonly employed in in vitro studies (for example, (Schnitzer et al., 2000)) and are within the ranges measured for intracellular cargos in vivo (Bakker et al., 1997; Casley-Smith, 1969; Keller et al., 2017; Lorenz and Willard, 1978; Margraves et al., 2011; Shubeita et al., 2008; Tytell et al., 1981; Wiemerslage and Lee, 2016; Zhang et al., 1998).

**Figure 1:**
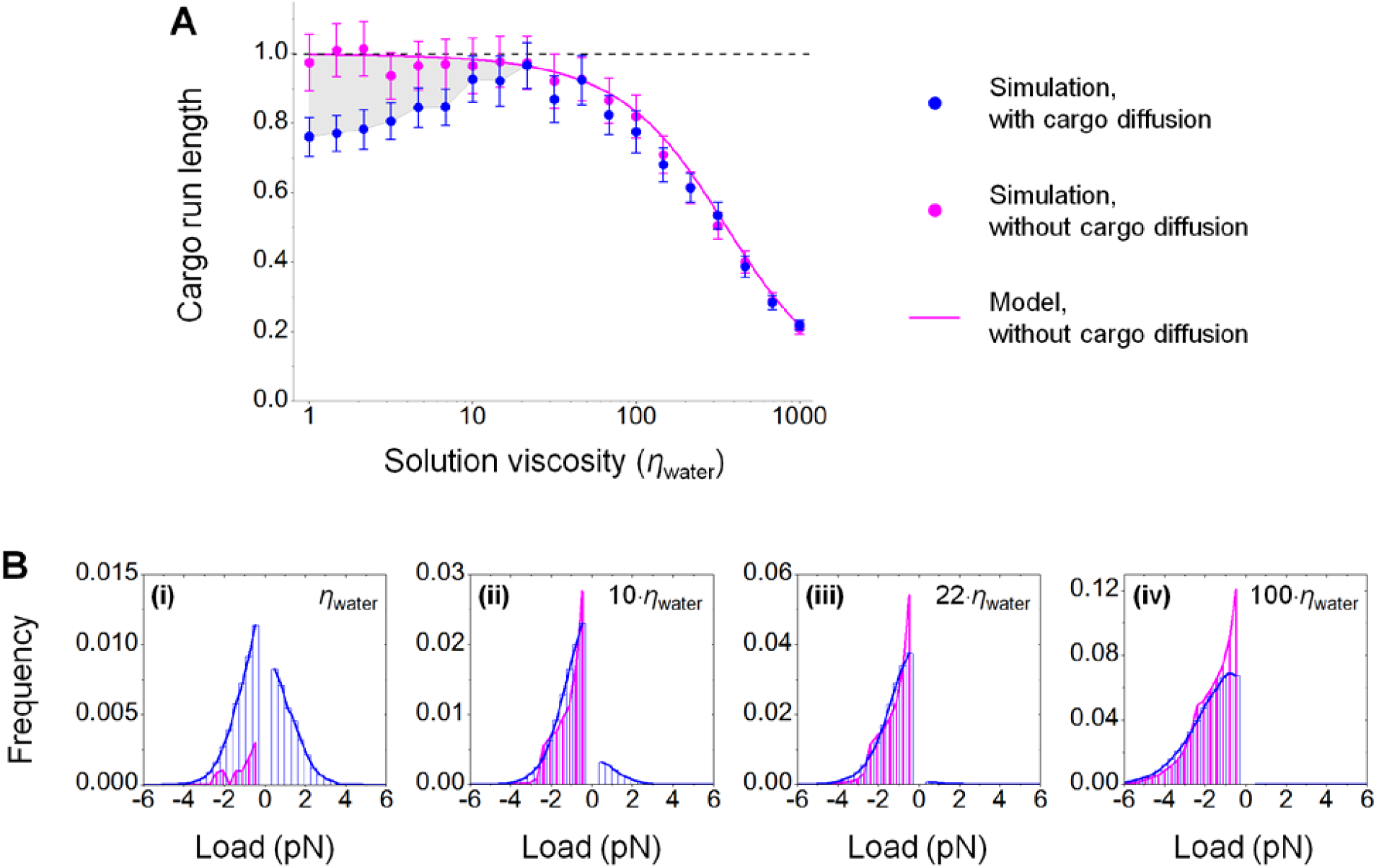
Thermal fluctuation of the cargo shortens single-kinesin run length at low viscosity (A) by imposing substantial assisting load on the motor (B). Simulations were carried out using a cargo 0.5 μm in diameter and a motor velocity of 0.8 μm/s unloaded. *η*_water_, the viscosity of water. (A) Grey area, the difference in run length between simulations with and without cargo diffusion. Error bars, standard error of the mean. (B) Positive values indicate load in the direction that assists motor movement; negative values indicate load in the direction that hinders motor movement.

Perhaps surprisingly, our simulations revealed a non-monotonic dependence of cargo run length on solution viscosity (blue scatters, Figure 1A). Whereas the mean cargo run length reached only 76±6% of the unloaded single-kinesin value in aqueous buffer, it increased significantly (*p* = 2×10^−4^, rank-sum test) to 97±7% of the unloaded single-kinesin value at a solution viscosity ~22-fold higher than that of the aqueous buffer, before declining with further increases in solution viscosity (blue scatter, Figure 1A). In contrast, when we did not include the thermal diffusion of the cargo in our simulations, we did not detect a non-monotonic dependence of cargo run length on solution viscosity (magenta scatter, Figure 1A). Our simulation of the diffusion-free case is in excellent agreement with predictions of the analytical model that considers the motor’s response to viscous load but not cargo diffusion ((Bell, 1978; Klumpp and Lipowsky, 2005; Singh et al., 2005), Methods) (magenta line, Figure 1A). We detected significantly shorter run length for simulations with cargo diffusion than the diffusion-free case (for example, *p* = 9×10^−4^, rank-sum test, for simulations at the viscosity of water) (grey area, Figure 1A). This difference in run length vanished at higher viscosities, where viscous drag alone was sufficient to shorten run length substantially (magenta, Figure 1A).

Together, our data demonstrate that thermal diffusion of the cargo shortens kinesin run length from that of the diffusion-free case. This effect is pronounced at low viscosity (grey area, Figure 1A), yielding a non-monotonic dependence of cargo run length on solution viscosity.

### Cargo diffusion imposes assisting load on the motor that is absent in the diffusion-free case

How does cargo diffusion shorten single-kinesin run length at low viscosity? Molecular motors such as kinesin are affected by mechanical load; the shorter run length suggests a greater load on the motor (Milic et al., 2014; Schnitzer et al., 2000). We thus hypothesized that cargo diffusion increases the load on the motor at low viscosity. To test this hypothesis, we compared the distribution of load on the motor during simulations with and without cargo diffusion (blue and magenta, Figure 1B).

We found that thermal diffusion of the cargo introduced substantial assisting load on the motor at low viscosity (positive load, blue, Figure 1B, i-iii). For example, at the viscosity of water, the motor had a similar probability of experiencing load in the assisting direction as in the hindering direction (blue, Figure 1Bi). In contrast, in the diffusion-free case, the motor experienced load only in the hindering direction (magenta, Figure 1Bi), which is expected because viscous drag always opposes cargo motion (Laidler and Meiser, 1982). Note that cargo diffusion also increased the hindering load on the motor at low viscosity (negative load, blue vs. magenta, Figure 1Bi). For example, at the viscosity of water, the motor had a higher probability of experiencing a greater hindering load in the presence of cargo diffusion than in the diffusion-free case (blue vs. magenta, Figure 1Bi). This observation is reasonable: thermal diffusion of the cargo is not correlated with the direction of motor motion (Einstein, 1905) and can thus contribute to load in both directions. However, as viscosity increased, the contribution of cargo diffusion to hindering load diminished more quickly than did the contribution of cargo diffusion to assisting load (Figure 1B, i and iii).

Taken together, our data demonstrate that cargo diffusion imposes substantial assisting load on the motor at low viscosity. Because assisting load *shortens* kinesin’s run length more severely than does hindering load (Milic et al., 2014), a substantial assisting load associated with cargo diffusion supports the observed reduction in run length versus the diffusion-free case.

### Effect of cargo diffusion on run length depends non-monotonically on viscous drag force

We next sought to understand how changes in cargo size and motor velocity impact kinesin’s run length. While these parameters were held constant in the preceding simulations at 0.5 μm in diameter and 0.8 μm/s unloaded, respectively (Figures 1), their values are known to vary in living cells (Bakker et al., 1997; Casley-Smith, 1969; Keller et al., 2017; Lorenz and Willard, 1978; Margraves et al., 2011; Shubeita et al., 2008; Tytell et al., 1981; Wiemerslage and Lee, 2016; Zhang et al., 1998).

We first examined the impact of cargo size, while holding motor velocity constant at 0.8 μm/s unloaded. The effect of solution viscosity on run length remained non-monotonic for cargos 0.1-1 μm in diameter (left, Figure 2A). Interestingly, the viscosity at which the run length of the single-kinesin cargo most closely approached the unloaded single-motor value (“critical viscosity”) scaled inversely with cargo size (dashed line, left, Figure 2A). The larger the cargo, the smaller the critical viscosity (dashed line, left, Figure 2A). This inverse scaling suggests that the effect of cargo size (*d*) and solution viscosity (*η*) on run length can be summarized as that of their product *dη* (Opper and Saad, 2001). Consistent with this hypothesis, the simulated run lengths for each combination of solution viscosity and cargo size collapsed onto a single curve with *dη* as the control parameter (left, Figure 2B).

**Figure 2.**
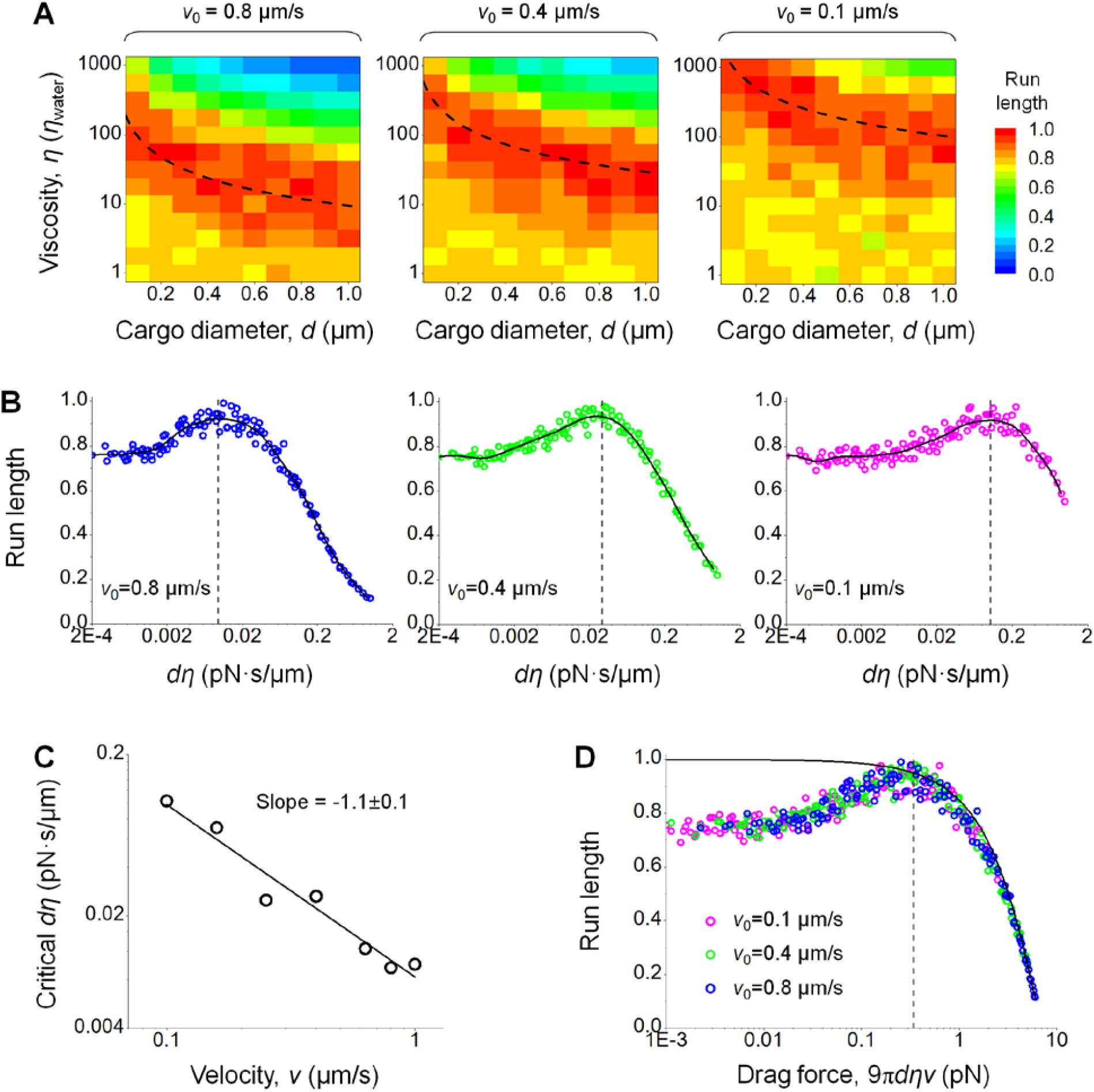
Non-monotonic variation in run length is general for physiologically relevant ranges of cargo size and motor velocity and is summarized by the single parameter of viscous drag force. *ν*_0_ indicates the unloaded motor velocity. (A) Dashed line, critical viscosity (where run length approached that of the unloaded single kinesin), which scales inversely with cargo size. *η*_water_, the viscosity of water. (B) For each unloaded motor velocity, the impact of cargo size and solution viscosity on run length (panel A) can be summarized as that of their product *dη*. Solid curve, moving average-smoothed data to guide the eye. Vertical dashed line, critical *dη* value, where run length approaches that of the unloaded single kinesin. (C) The critical *dη* value scales inversely with motor velocity. Solid line, best linear fit with the indicated slope. (D) The impact of *dη* and motor velocity on run length (panel B) can be summarized as that of viscous drag force. Solid line, model predictions of cargo run length as a function of viscous drag force. Vertical dashed line, an approximate threshold where the effect of viscous drag force on cargo run length becomes substantial.

We next examined the impact of motor velocity on our simulation results (Figure 2, A-C). For each unloaded motor velocity examined, the run lengths of single-kinesin cargo again varied non-monotonically with the combined parameter *dη* (for example, 0.4 μm/s and 0.1 μm/s, Figure 2, A and B). Interestingly, the value of *dη* at which cargo run length approached the unloaded single-motor run length correlated inversely with motor velocity (Figure 2C). This inverse scaling suggests that the effects of *dη* and motor velocity (*ν*) on run length may be again combined as that of their product *dην* (Opper and Saad, 2001), or the drag force experienced by the cargo (9*πdην*, (Chen and Ye, 2000; Laidler and Meiser, 1982)). Consistent with this suggestion, the run length for each of the three unloaded motor velocities (Figure 2C) collapsed onto a single curve with the drag force as the single control parameter (Figure 2D).

Thus, the run length of single-kinesin cargos in a viscous medium is influenced by three independent parameters: solution viscosity, cargo size, and motor velocity. The effect of these three parameters on cargo run length can be summarized as that of a single control parameter: the product of the three parameters, or the viscous drag force that arises from the active motion of the motor. This collapsed single-parameter curve differs substantially from model predictions for the diffusion-free case at low viscous drag forces; this difference diminishes when the effect of viscous drag force on kinesin’s run length becomes pronounced (scatters vs. solid line, Figure 2D).

### Viscous drag force biases thermal diffusion of cargo toward the hindering direction

How does viscous drag force—arising from motor motility—alter the effect of cargo diffusion on kinesin run length? To address this question, we examined the displacement of the diffusing cargo diffusion from the motor, which determines the load that the diffusing cargo exerts on the motor, for a range of viscous drag forces (Figure 3).

**Figure 3.**
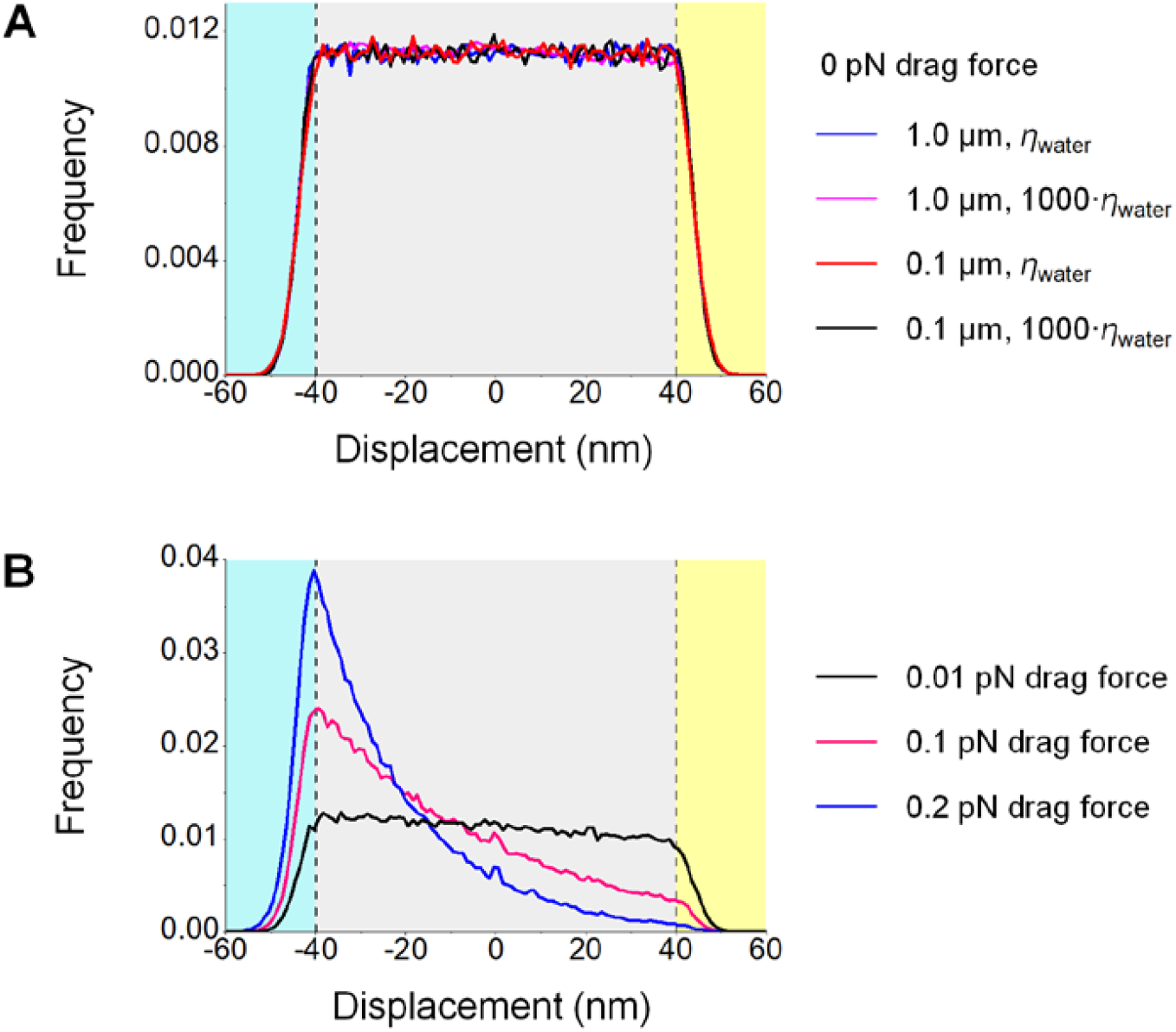
The distribution of displacement of a thermally diffusing cargo from the motor is sensitive to viscous drag force. Simulations were carried out at zero drag force (A) and at low drag forces (B). Positive displacement reflects the cargo leading in front of the motor; negative displacement indicates that the cargo lags behind the motor. Grey area, free-diffusion range where the cargo does not impose any load on the motor. Cyan (and yellow) area, tethered-diffusion range where the cargo imposes hindering (and assisting) load on the motor. (A) At zero drag force, the displacement of a diffusing cargo is symmetric about the motor position (0 nm) and is not sensitive to cargo size or solution viscosity (*p*=0.70, Kruskal–Wallis one-way analysis of variance). (B) At non-zero drag forces, the displacement of a diffusing cargo is biased toward the hindering direction (negative displacement). The magnitude of this bias increases as the viscous drag force increases.

We first carried out simulations for the case of zero drag force (Figure 3A). Here, the motor velocity was kept constant at 0 μm/s, and solution viscosity and cargo size were varied over the physiologically relevant range used in preceding simulations (1000-fold and 10-fold, respectively). The resulting displacement distributions were symmetric about the motor position and exhibited two diffusion regimes: a uniformly distributed “free diffusion” range (grey area, Figure 3) where thermal motion of the cargo does not stretch the motor beyond its natural length and is thus effectively decoupled from the motor (Einstein, 1905), and a normally distributed “tethered diffusion” range (cyan and yellow areas, Figure 4) where thermal excursion of the cargo is restricted by the motor that tethers the cargo to the microtubule (Pathria and Beale, 2011). Characteristics of these two diffusion regimes were not significantly influenced by cargo size or solution viscosity (*p*=0.70, Kruskal–Wallis one-way analysis of variance, Figure 3A). Each simulated displacement distribution demonstrated a similar frequency of occurrence within the free-diffusion range (~0.01, grey area, Figure 3A) and a similar variation in the tethered diffusion range (~7.5 nm standard deviation in both load directions, cyan and yellow areas, Figure 3A). These displacement distributions correspond to a 30% increase in the motor’s detachment rate and a 26% reduction in motor run length from their unloaded values. These observations are in excellent agreement with the ~24% reduction in run length in our simulations at negligible viscous drag force (1×10^−3^ pN, Figure 2D).

**Figure 4.**
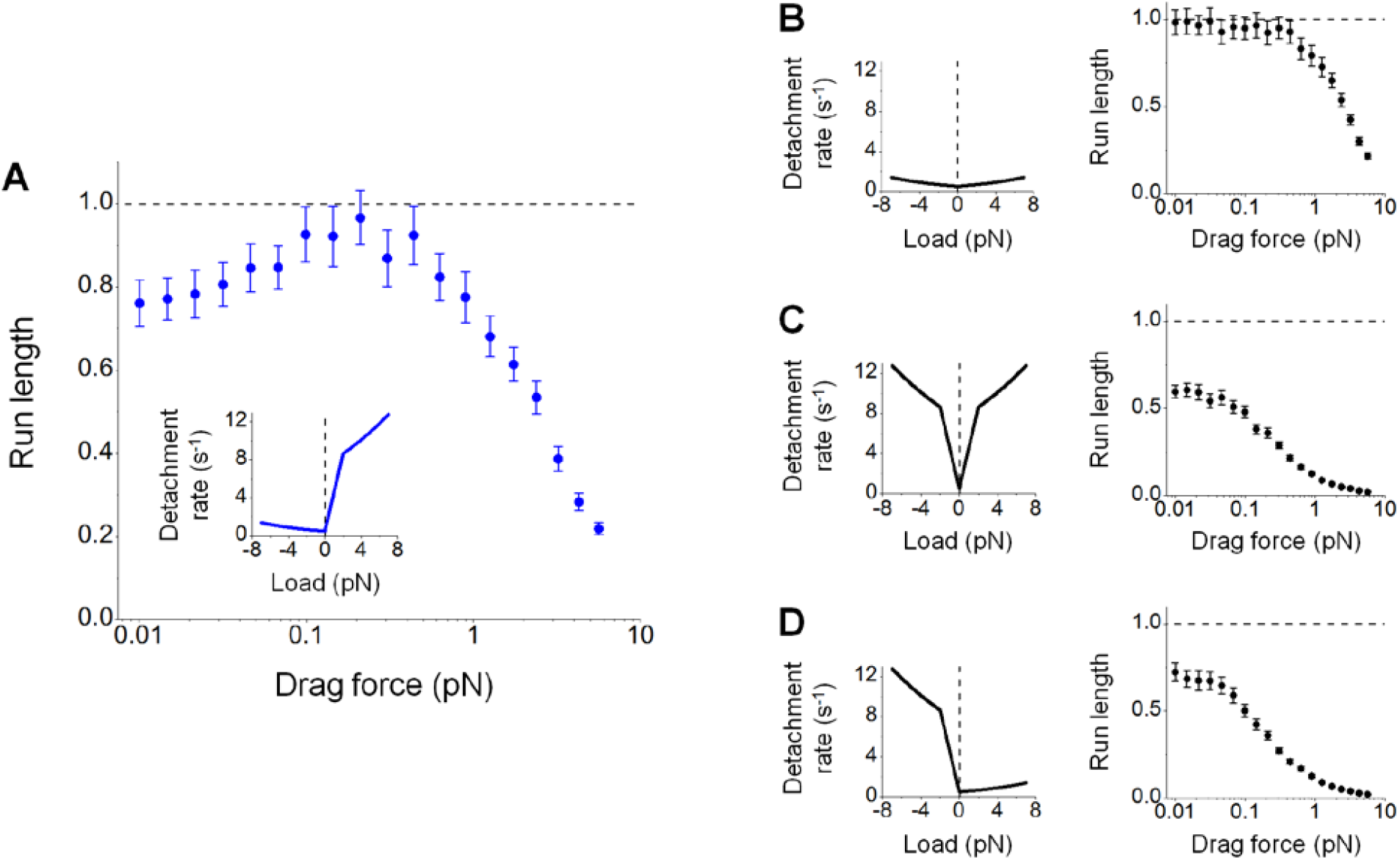
Non-monotonic dependence of cargo run length on viscous drag force requires a specific symmetry of the motor’s load-detachment kinetics. Error bar, standard error of the mean. (A) Experimentally measured load-detachment kinetics (Milic et al., 2014) (inset) give rise to a non-monotonic dependence of cargo run length on viscous drag force. Data here are duplicated from Figure 1A (blue scatter). (B-C) Symmetric load-detachment kinetics (left) cannot support a non-monotonic dependence of cargo run length on viscous drag force (right). (D) Asymmetric load-detachment kinetics with reduced sensitivity for assisting versus hindering load (left) also cannot support a non-monotonic dependence of cargo run length on viscous drag force (right).

We next examined the case of non-zero viscous drag force (Figure 3B). Here, the motor velocity was kept constant at 0.8 μm/s, and solution viscosity and cargo size were chosen to capture the low drag force range that alleviates the shortening effect of cargo diffusion on kinesin run length (0.01-0.2 pN, Figure 2D). The resulting displacement distributions were asymmetric about the motor position in both the free-diffusion range (grey area, Figure 3B) and the tethered-diffusion range (cyan and yellow areas, Figure 3B). As the viscous drag force increased, the position of the diffusing cargo increasingly lagged behind the motor. At a drag force of 0.2 pN, the position of the cargo exhibited a <0.4% probability of leading the motor position (blue line, Figure 3B). Of note, despite the asymmetry in the displacement distribution, variations in the tethered diffusion range remained similar between load directions (~7.6 nm standard deviation in the hindering direction and ~7.5 nm standard deviation in the assisting direction, cyan and yellow areas, Figure 3B) and unchanged from that for the zero drag case (~7.5 nm in both load directions, cyan and yellow areas, Figure 3A).

Together, our simulations indicate that viscous drag biases the diffusing cargo to lag behind the moving motor, thereby reducing the likelihood of the motor experiencing assisting load. At low viscous drag force, this reduction in assisting load is accompanied by an increased likelihood, but not magnitude, of hindering load on the motor.

### Non-monotonic tuning by viscous drag force requires specific asymmetry in the motor’s load-detachment kinetics

We next sought to understand how the motor’s load-detachment kinetics impacted our simulation results. We hypothesized that non-monotonic tuning by viscous drag force (Figure 2D) reflects a selective sensitivity of the motor’s detachment rate to the combination of lower assisting load and higher hindering load (Figure 3B). To test this hypothesis, we varied the symmetry properties of the motor’s load-detachment kinetics under otherwise identical simulation conditions (Figure 4). For ease of comparison, we duplicated our preceding simulations using experimentally measured load-detachment kinetics for single kinesins (Milic et al., 2014).

We found that asymmetry in kinesin’s load-detachment kinetics is necessary but not sufficient for the observed non-monotonic dependence of cargo run length on viscous drag force (Figure 4). We first examined the effect of symmetric load-detachment profiles on cargo run length (Figure 4, B and C). Here, we duplicated the experimentally measured load dependence (Milic et al., 2014) in the hindering direction (left, Figure 4B) or the assisting direction (left, Figure 4C). In both cases, the effect of drag force on cargo run length increased monotonically, with cargo run length maintaining its maximum value at the lowest drag force tested (right, Figure 4, B and C). As expected, the maximum run length of the single-motor cargo was substantially lower when we assumed a higher sensitivity of the motor’s unbinding rate to load (right, Figure 4, C vs. B). We next generated an asymmetric load-detachment profile that preserved the magnitude of impact but reversed the directional bias of the measured load dependence (left, Figure 4D); this assumed load-detachment profile again yielded a monotonic dependence of cargo run length on viscous drag force (right, Figure 4D).

Hence, these simulations reveal that the specific asymmetry in the recently measured load-detachment kinetics of kinesin—greater sensitivity of the motor’s detachment rate to assisting load than to hindering load (Milic et al., 2014) (Figure 4A, inset)—underlies the non-monotonic dependence of cargo run length on viscous drag force. An interesting implication of this finding is that the effect of viscous force on cargo velocity should be monotonic, because the load-stepping kinetics of kinesin does not satisfy the specific asymmetry identified here (Milic et al., 2014; Schnitzer et al., 2000) (inset, Figure 5). Velocities associated with the simulated run lengths in Figures 4A support this implication (Figure 5).

**Figure 5.**
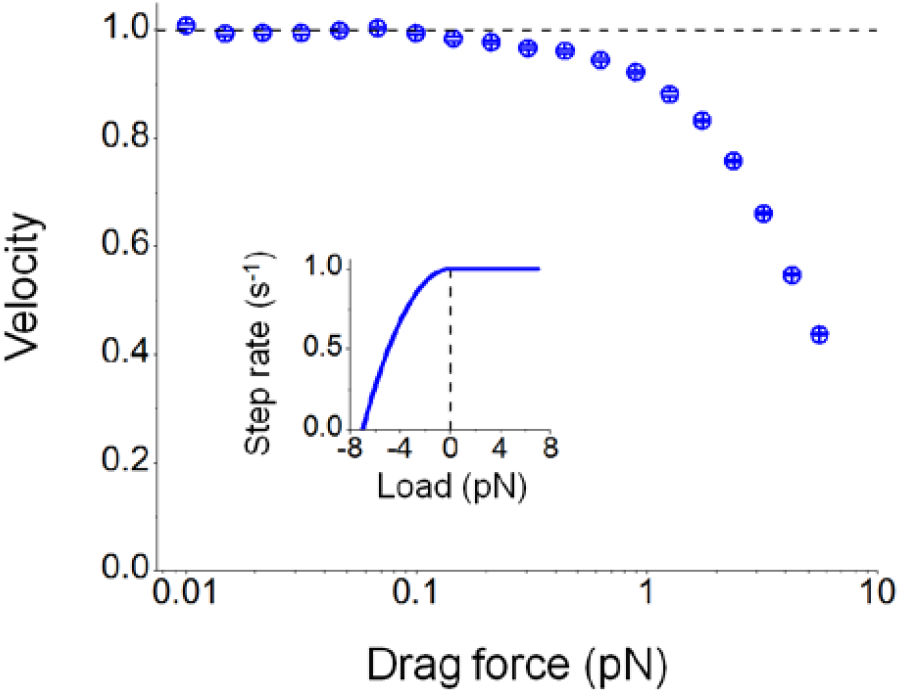
The velocity of single-kinesin cargos decreases monotonically as viscosity increases. Velocities were evaluated for simulations used in Figures 4A. Error bar, standard error of the mean. Inset, load-stepping kinetics of the motor (Milic et al., 2014; Schnitzer et al., 2000) does not satisfy the specific asymmetry requirement identified in the current study (inset, Figure 4A) and does not support a non-monotonic dependence of cargo velocity on viscous drag force.

## Discussion

Here we employed Monte Carlo-based stochastic simulations to examine the effect of thermal diffusion of the cargo impacts the run length of single-kinesin cargos. Our simulations revealed that cargo diffusion shortens single-kinesin run length, by imposing substantial assisting load on the motor that is absent in the diffusion-free case. Viscous drag force biases the effect of cargo diffusion toward hindering load. This interplay between viscous drag force and cargo diffusion, combined with the asymmetric response of kinesin’s run length to load direction, gives rise to a previously unexplored, non-monotonic dependence of cargo run length on viscous drag force.

Our study highlights the importance of load-detachment kinetics on the functions of single motor carrying a thermally diffusing cargo. The shortening effect of cargo diffusion on kinesin run length uncovered here arises from the motor’s sensitivity to the thermally based assisting load. The more likely that the motor detaches under load in the assisting versus the hindering direction, the greater the shortening effect of cargo diffusion on the motor’s run length, and the greater the non-monotonic tuning of cargo run length by viscous drag. Although we focused our current study on the major microtubule-based motor kinesin-1, we anticipate similar shortening effects of cargo diffusion for other classes of motors that also demonstrate an enhanced sensitivity to assisting load, such as kinesin-2 (Andreasson, 2013) and cytoplasmic dynein (Nicholas et al., 2015), but not kinesins-3, 5, or 7 (Arpag et al., 2014). In this respect, the load-detachment kinetics is likely a critical factor underlying diversity in the single-motor functions of distinct classes of motors.

Our characterization of viscous drag force as the single tuning factor for cargo run length may be important for understanding the diverse characteristics of transport observed in vivo. Because viscous drag force is determined by the product of three independent parameters (solution viscosity, cargo size, and transport velocity (Laidler and Meiser, 1982)), distinct conditions can give rise to the same magnitude of viscous drag force and hence the same run length. For example, the effect of a substantial, 100-fold reduction in solution viscosity may be countered by a 10-fold increase in cargo size and a 10-fold increase in cargo velocity; these changes are within physiological ranges.

In addition to the viscous effects examined here, the elastic nature of cytoplasm, such as that associated with spatial heterogeneity of the cytoskeleton (Katrukha et al., 2017), has been previously predicted to impact kinesin-based transport (Goychuk et al., 2014; Nam and Epureanu, 2012). Thermal diffusion of the cargo was not included in these important previous investigations. Future investigations combining solution viscoelasticity with cargo diffusion may reveal additional diversity or tunability in motor-based transport.

Finally, we focused the current study on cargo transport by a single kinesin-1. Intracellular cargos are often carried by small teams of motors (Hendricks et al., 2010; Kulkarni et al., 2017; Shubeita et al., 2008). Because transport by a small team of kinesin-1 molecules is on average accomplished by the action of a single kinesin (Furuta et al., 2013; Jamison et al., 2010; Xu et al., 2012b), our findings at the single-molecule level are likely directly relevant for interpreting the effect of cargo diffusion on transport by multiple kinesins. It will be interesting to examine the effect of cargo diffusion on team-motor transport, particularly for mixed motor populations. Recent investigations have focused on the importance of inter-motor strain (Belyy et al., 2016; Norris et al., 2014; Rogers et al., 2009) and local confinement (Feng et al., 2018; Xu et al., 2012b) on team-motor functions. The current study suggests an intriguing new consideration of how cargo exerts the thermally-based load on individual motors during team transport. We are developing simulations to address this question.

## Methods

### Monte Carlo-based stochastic simulation

A previously developed Monte Carlo-based stochastic simulation model (Kunwar et al., 2008) was adapted to evaluate the motility of single-kinesin cargos in a viscous medium. The current study used the numerical algorithm developed previously (Kunwar et al., 2008), but updated the motor’s load-detachment kinetics to reflect new experimental finding on the motor’s response to assisting load (Milic et al., 2014) as well as the previously known response to hindering load (Schnitzer et al., 2000).

Briefly, each cargo is carried by one motor, moving along a one-dimensional microtubule lattice. The motor is assumed to be an idealized spring with a native length and a linkage stiffness. The motor is assumed to exert a restoring force on the cargo only when it is stretched beyond its native length; the motor is assumed to have no compressional rigidity and buckles without resistance when compressed. A simulated cargo run is initiated when the motor becomes stochastically bound to the microtubule (characterized by the motor’s binding rate). At each simulation time step, the displacement between the cargo and the motor is used to determine the load on the motor (and on the cargo). This load information is used to determine the probability that the motor will become detached from the microtubule (characterized by the motor’s load-detachment kinetics). If the motor remains engaged in transport, then the load on the motor is used to calculate the probability that the motor will advance one step along the microtubule lattice (characterized by the motor’s load-stepping kinetics), and the position of the motor is updated accordingly. The position of the cargo at each time step is determined by summing the random thermal diffusion of the cargo and the deterministic drift of the cargo under load. The simulation time step is incremented and the above evaluations repeated until the motor stochastically detaches from the microtubule.

The simulation time step is chosen to range between 10^−6^ s and 10^−5^ s; no significant differences in simulation results were detected over this range of time steps (data not shown). Unless otherwise indicated, the same motor length (40 nm, (Kerssemakers et al., 2006)), motor stiffness (0.32 pN/nm, (Coppin et al., 1997)), binding rate (5/s, (Leduc et al., 2004)), step size (8 nm, (Coy et al., 1999; Schnitzer and Block, 1997)), stall force (7 pN, (Milic et al., 2014)), and unloaded run length (1.5 μm, (Xu et al., 2012a)) were used for all simulations. The values of solution viscosity, cargo size, and unloaded motor velocity are as indicated. The motor’s load-stepping kinetics was as determined in previous experimental investigations (Milic et al., 2014; Schnitzer et al., 2000). The motor’s load-detachment kinetics was as determined in the recent single-molecule experimental investigation (Milic et al., 2014). Specifically, the motor’s detachment rate under load *ε*(*F*) was described as the piece-wise function

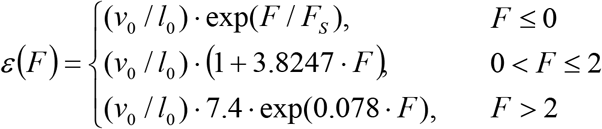

where *ν*_0_ is kinesin’s unloaded velocity, *l*_0_ is kinesin’s unloaded run length, *F_s_* is the single-kinesin stall force, and *F* is the load on the motor (in units of pN). Positive force indicates assisting load, negative force indicates hindering load.

### Data analysis

The run length of a simulated trajectory is defined as the overall distance traveled by the simulated motor before the cargo during transport. For each simulation condition, the cumulative probability distribution of the run lengths was fitted to the cumulative probability function of a single exponential distribution, 1 − *A* · exp(−*x*/*l*). Mean run length was determined as the best-fit decay constant *l*. The associated standard error of the mean was determined via a bootstrap method (Manly, 2006). Mean run lengths in all figures were normalized by the unloaded single-kinesin run length.

The velocity of a simulated trajectory was determined as the best-fit slope of the trajectory. Only trajectories with ≥0.1 μm of motion and ≥0.2 s of duration were analyzed. For each simulation condition, mean velocity was calculated as the arithmetic mean. The associated standard error of the mean was determined by a bootstrap method (Manly, 2006). Mean velocities in Figure 5 were normalized by the unloaded single-kinesin velocity.

The load on the motor was determined using the displacement of the cargo from the motor during each simulated run. Displacements shorter than the native length of the motor do not impose any load on the motor. Displacements longer than the native length of the motor impose a load on the motor. The magnitude of the load was determined as the length of the motor stretched from its native value, multiplied by motor stiffness. The direction of the load was determined by the relative position of the cargo to the motor: “assisting” when the cargo position leads the motor, “hindering” when the cargo position lags behind the motor.

The detachment rate of the cargo for a given displacement distribution in Figure 4 was determined as the weighted sum of the kinesin’s detachment rate at a particular displacement value (corresponding load evaluated as described above), multiplied by the frequency of that particular displacement value as the weight. Kinesin’s detachment rate at a particular displacement value was calculated by first determining the load associated with the displacement value, then applying the motor’s load-detachment kinetics as described above. The associated run length of the cargo was determined as the ratio of the cargo’s velocity to its detachment rate.

### Statistical analysis

The rank-sum test was used to detect significant differences between two distributions of cargo run lengths. Kruskal-Wallis one-way analysis of variance was used to evaluate the significance of differences between multiple distributions of cargo displacement in Figure 4.

### Analytical model of kinesin run length in the absence of cargo diffusion

In the absence of cargo diffusion, the only load on the motor is imposed by the viscous drag force in the direction hindering the motor’s motion. Based on the experimentally measured load-detachment kinetics of kinesin for hindering loads (Milic et al., 2014; Schnitzer et al., 2000), the run length of single-kinesin cargos in viscous medium (*l*) is modeled by the Bell equation (Bell, 1978; Klumpp and Lipowsky, 2005; Singh et al., 2005):

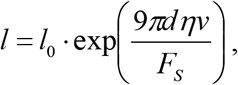

where *l*_0_ is the unloaded single-kinesin run length, *F_s_* is the single-kinesin stall force, *d* is the cargo diameter, *η* is the solution viscosity, and *v* is the velocity of the motor under viscous load. Note that the viscous drag force on the cargo, 9π*dην*, includes consideration of the ~3-fold higher viscosity near the surface than in the bulk solution, per Faxen’s law (Chen and Ye, 2000).

The loaded velocity of the motor in the preceding equation was calculated as follows. Based on the experimentally measured load-stepping kinetics of kinesin (Milic et al., 2014; Schnitzer et al., 2000), the velocity of the motor under viscous load satisfies the quadratic relationship

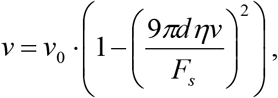

where *ν*_0_ is the unloaded single-kinesin velocity, and 9*πdην* is the drag force on the cargo as described above. The solution to this quadratic equation gives rise to the analytic description of the velocity of single-kinesin cargos in a viscous medium

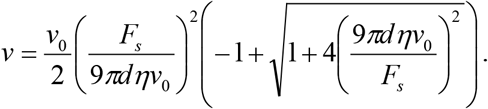

## Acknowledgments

We thank Bayana Science for manuscript editing.

We acknowledge support from the National Institutes of Health (R15 GM120682 to J.X.). J.O.W. acknowledges partial support from the National Science Foundation—CREST: Center for Cellular and Biomolecular Machines at UC Merced (No. NSFHRD-1547848). Numerical simulations in this study were carried out using the Multi-Environment Research Computer for Discovery cluster computing resource supported by the National Science Foundation (ACI-1429783).

